# Intriguing density dependence but problematic birth rates in Greaver et al.’s study of a black-rhinoceros population

**DOI:** 10.1101/039453

**Authors:** Peter R Law, Wayne L. Linklater, Jay V. Gedir

## Abstract

Greaver et al. (2014) presented evidence for density dependence in the Ithala population of black rhinoceros. Finding that they did not place their regression-based evidence in a modelling context, we recast their result as an example of the ramp model of density dependence that underlies black rhinoceros meta-population management. Greaver et al. concluded that the Ithala population did not reach carrying capacity, a conclusion we consider unwarranted since they did not conduct any analyses of trends in demographic parameters with population density. Our interpretation implies that the Ithala population did indeed reach carrying capacity. Where relevant, we compared their results for the Ithala population with those for another southern African black-rhinoceros population in order to provide a broader basis for evaluating black rhinoceros demography. We detail inconsistent presentation of data in their paper that plagued our efforts to understand their results and also draw attention to possible errors in some analyses. In particular, we argue that the results on birth rates reported by Greaver et al. appear dubious. Greaver et al. have presented important evidence for density dependence in a population of black rhinoceros but we suggest they have underutilized their data in interpreting this density dependence while misanalysing birth data.

## Introduction

Current understanding of black rhinoceros (*Diceros bicornis*) demography and population dynamics remains poor. Detailed demographic datasets spanning even rhinoceros’ lifetimes, let alone multiple generations, would be challenging enough to compile in even ideal conditions but are all the scarcer due to the considerable disturbances their populations have suffered through poaching and habitat loss. It is only recently that studies of black-rhinoceros demography have progressed much beyond natural history. Notable examples include Cromsigt et al. (2002), Hrabar and du Toit (2005), Okita-Ouma et al. (2009), Brodie et al. (2011), Ferreira et al. (2011), and Law et al. (2013, 2014, 2015).

It was therefore with considerable interest that we greeted the publication of Greaver et al. (2014), who had access to 19 years of quality data on a population of black rhinoceros (Ithala Game Reserve, South Africa). While 19 years of data is still limited for a species whose individuals can live into their thirties, it represents sustained and careful monitoring by dedicated staff that should be capitalized on to increase our understanding and inform conservation policy and practice.

An important topic for understanding the ecology of rhinoceros that still requires elucidation is the form in which density dependence acts on their populations (and those of megaherbivores in general) and how density dependence is mediated. This aspect of rhino ecology is vital because many populations of rhinoceros are now confined by fences on reserves and require intensive management to avoid overpopulation of reserves and maintenance and expansion of the meta-population through translocations. Meta-population management of black rhinoceros in Africa has been based (Emslie 2001) on a so-called ramp model in which annual per capita growth rate is constant until the population size nears carrying capacity and then declines rapidly (Fowler 1981, 1987; McCullough 1992, 1999). Determining whether the ramp model is in fact appropriate and, even if it is, what the demographic indicators of the approach to carrying capacity are, requires quality data collected over many years and is thus an important issue that Greaver et al. might address.

Indeed, their paper contains a regression, their Fig. 2b, that relates to this very question. Reflexion upon their regression leads us to propose an interpretation of the regression consistent with a ramp model. While both we and Greaver et al. concur the regression is evidence for density dependence, we think they have dismissed the possibility that carrying capacity was reached on inadequate grounds and left the Ithala data underutilized, with the possibility that their conclusions as to the character of density dependence might be misleading.

While drawing attention to this very interesting aspect of their paper, we are compelled to point out difficulties we encountered in comprehending the data as presented by Greaver et al. These difficulties led us to scrutinize the paper carefully, which raised further questions about some of the analyses and conclusions, which we also wish to address here. In addition to inconsistent presentation of data, which plagued our attempts to reconcile several figures and results, we contend that their analyses and results on birth rates appear dubious.

We will also take this opportunity to make some comparisons between the Ithala black-rhinoceros population and another black rhinoceros population in the Great Fish River Reserve, South Africa, the demographics of which were studied in Law et al. (2013, 2014, 2015). As in those publications, we will refer to this population as the SKKR population. It was founded through reintroduction, beginning in 1986, with the last release in 1997. The study treated the years 1986 through 2008. The SKKR population grew exponentially during the study period with no discernible density dependence in growth (Law et al. 2105). Thus, the SKKR dynamics were different to those of the Ithala population as reported by Greaver et al. Comparisons between these two populations highlight interesting and informative variation in the dynamics of reintroduced populations of black rhinoceros but also supports some of our concerns about the results reported by Greaver et al.

### Density Dependence and Population Growth Rate

We begin by considering the very interesting regression in Greaver et al.’s Fig. 2b, which they presented as evidence for density dependence. We have reproduced their Figures 2 and 4, with annotations, as our Figures 1 and 2.

**Figure 1.**
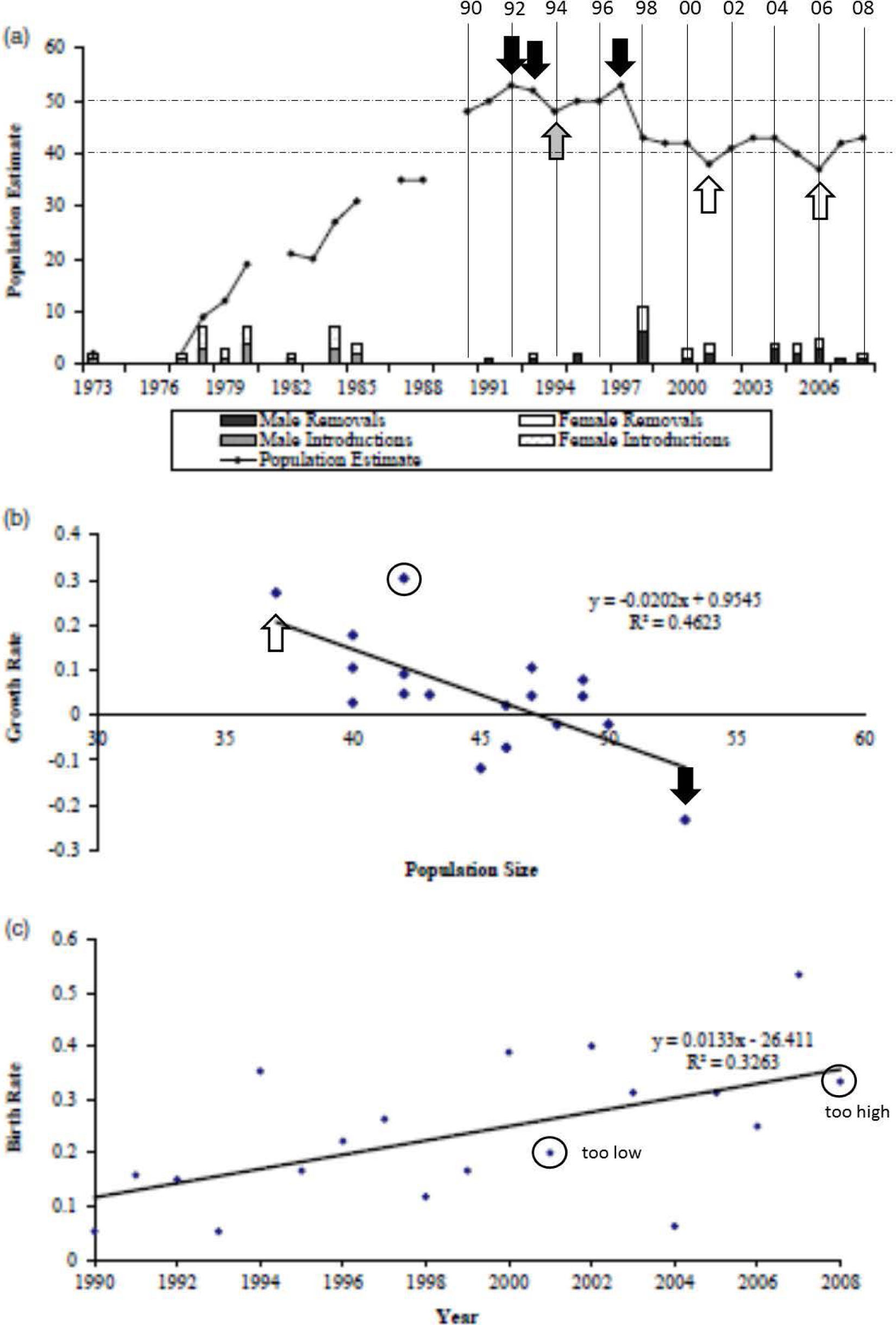
Annotated copy of Fig. 2a-c from Greaver et al. (2014) to illustrate points made in the text.

**Fig. 2.**
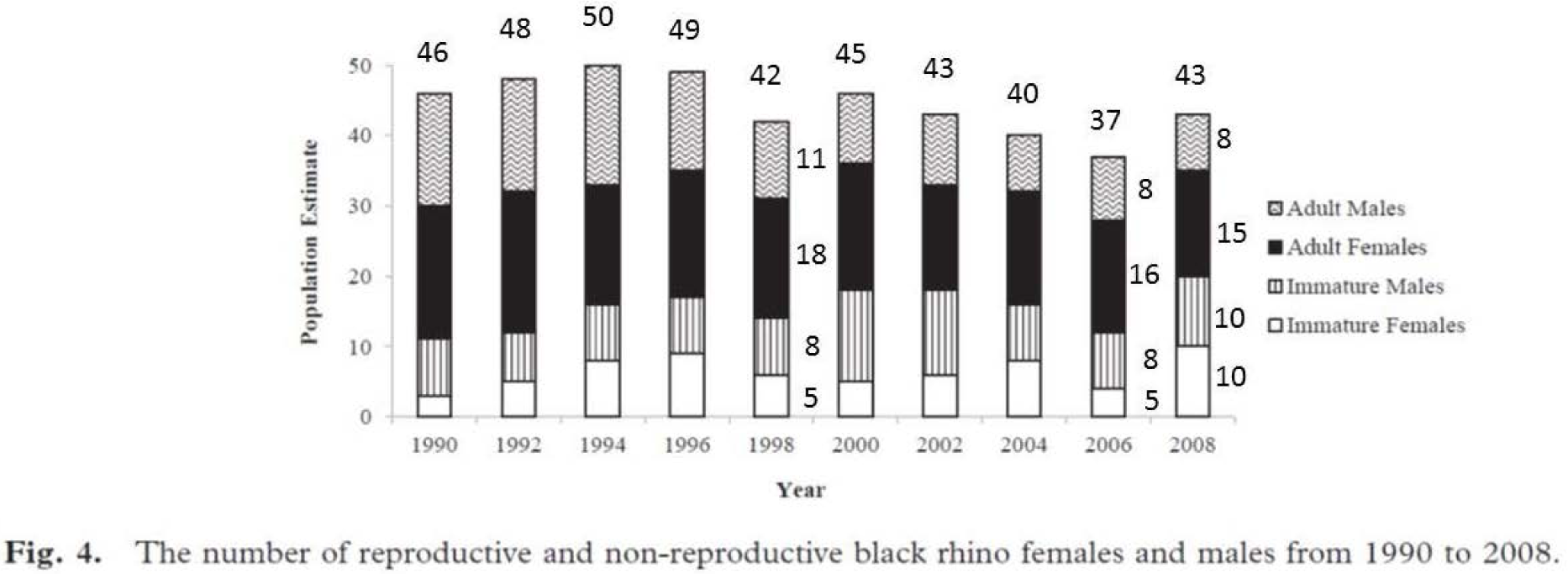
Black rhino population (a) observed estimates since the introduction in Ithala in 1973 (b) growth rates (c) birth rates from 1990 to 2008. (Greaver et al.’s caption to their Figure 2) (a) We use solid, labelled vertical lines to clarify the match between data points and bars, and year of the study (the x-axis labels in the original figure do not align with the data). The horizontal, dashed lines indicate population sizes of 40 and 50. The arrows indicate three data points above 50 (black), one just below 50 (grey), and two below 40 (open). (b) The open arrow indicates the only population size less than 40, at ~(37,0.27), although two are indicated in Fig. 2a. The black arrow indicates the only data point at a population size larger than 50 although three such data points are shown in Fig. 2a. The circled data point, at ~(42, 0.29), has the highest r value and is the point most deviant from the regression and is also a test point for our interpretation of the data. (c) The circles indicate data points whose birth rates we could not reproduce regardless of how we interpreted the data, see text. (Law et al.’s caption to our Figure 1) Fig. 4 from Greaver et al. (2014) annotated to show our interpretation of the sizes of histogram bars.

We first make an observation about the values that do not fit the regression well. The higher positive values of *r* correspond to years in which a high proportion of adult females calved, while the negative value of high magnitude must reflect high mortality. The latter value of *r* does occur for the highest abundance. For comparison, in SKKR, the two highest values of *r* occurred for 1993, with an initial population of 15 and 4 births during the year, and 2000, with a population of 46 and 11 births during the year. Thus, some years of exceptional growth, not conforming to the regression, are not surprising. Figure 3 provides a plot of the *r* values for SKKR, computed as in Greaver et al. (p. 439) by subtracting any removals/additions during the year from the starting/ending population size and then taking the logarithm of the ratio of the modified ending and starting population sizes. Linear regression returned *F*_1,19_ = 0.044, *p* = 0.836, *R*^2^ = 0.061, *R*^2^_a_ = −0.050, the intercept was 0.095, the slope 0.0005. In other words, regression does no better than the mean. Indeed, amongst scalar models of population growth based on the generalized logistic, AIC_c_ unambiguously selected a model of exponential growth for SKKR (Law et al. 2015), with no evidence of density dependence in growth rates.

**Figure 3.**
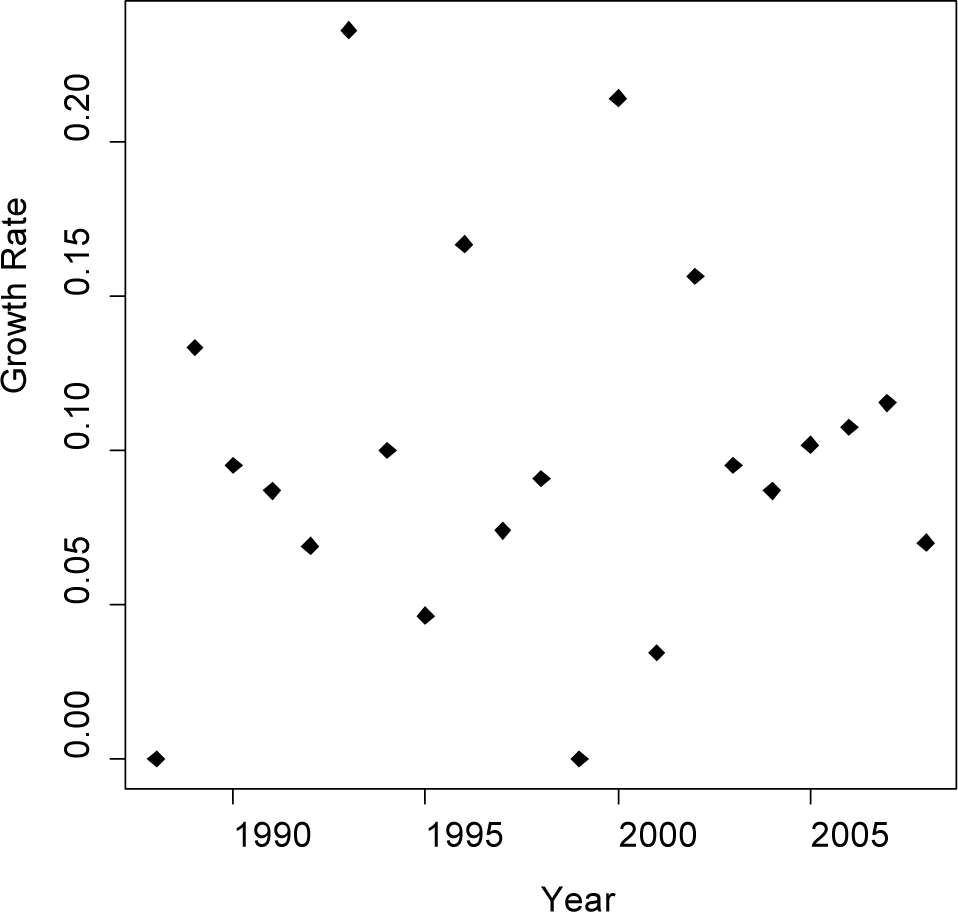
Values of *r* for the SKKR population of Law et al. (2013, 2014, 2015).

A linear regression of the form *r* = a + b*N*_*t*_ as in Greaver et al.’s Fig. 2b can be interpreted as

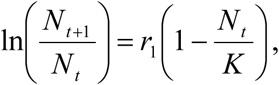

which amounts to fitting the Ricker model (Owen-Smith 2010:38) to a time series of population counts. Doing so assigns values of *K* = 47.25 (the value where the regression crosses the horizontal axis) and *r*_1_ = 0.955 (Greaver et al. did not provide SEs for their regression coefficients). Normally, *r*_1_ would be interpreted as the intrinsic rate of growth for a population growing according to the Ricker model. Estimates of the intrinsic rate of growth of black rhinoceros populations reported in the literature are typically about 0.1 or smaller (Law et al. 2015). The value 0.955 is therefore unrealistically high for the intrinsic rate of growth, indicating that the regression does not extrapolate to smaller population sizes beyond the range of data on which it was based.

Unfortunately, it appears the Ithala data prior to 1990 is not good enough to model. Nevertheless, the ramp model of density dependence suggests patching exponential growth to the above Ricker model. If *r*_0_ denotes the intrinsic rate of growth for the population, then one requires

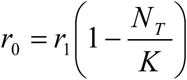

at a threshold population size *N*_*T*_ at which exponential growth ceases and the Ricker model comes into effect. Greaver et al.’s regression determines *r*_1_ and *K*, so *r*_0_ and *N*_*T*_ determine each other by this equation. In the absence of reliable data for the population dynamics prior to 1990, if we take *r*_0_ ≈ 0.1, based on the literature, then *N*_*T*_ ≈ 42.3 ≈ 90% of *K* (the threshold size is the abundance at which the horizontal line at height 0.1 crosses the regression line). Note that lower intrinsic rates of growth yield higher threshold values *N*_*T*_. Thus, one possible interpretation of Greaver et al.’s Fig. 2b is that there were a couple of years of high growth rate (due to some stochasticity) but that basically we are observing the declining part of a ramp model of population dynamics. It is unfortunate that the Ithala data prior to 1990 is not good enough to test our proposed ramp model during the growth phase of the Ithala population, which might thereby have provided direct evidence for the full ramp model of density dependence. Other studies, however, have reported sustained exponential growth in populations of black rhinoceros (Okita-Ouma et al. 2010, Law et al. 2015) while Cromsigt et al. (2002) presented evidence for ramp-like density dependence. Our interpretation of Greaver et al.’s regression is an intriguing possibility that is consistent with theoretical expectations of megaherbivore population dynamics (Fowler 1981, 1987; McCullough 1992, 1999), which thereby provides an ecological context for the regression. Our interpretation is at odds with that of Greaver et al., however, as we next discuss.

### Social and/or resource mediation of density dependence?

Greaver et al. asserted (p. 445) that the Ithala population had not reached ecological carrying capacity, in the sense of not being resource limited, because the mean age at first reproduction (AFR) and mean inter-birthing interval (IBI) they reported do not suggest so. Their mean AFR was 6.5 (SE = 0.42) years (*n* = 18), i.e., 78 months (SE= 5.04) not dissimilar to the 80 (SE = 3.7) months (*n* = 16) of the rapidly growing SKKR population. Their mean IBI was 3.2 (SE = 0.04) years, i.e., 38.4 (SE = 0.48) months (*n* = 61) while for SKKR it was 29.0 (SE = 0.9) months (*n* = 77). Thus, while their mean AFR was very similar to that for SKKR, their mean IBI was substantially longer. Moreover, Greaver et al. did not study trends in either AFR or IBI with population size or, for example, rainfall (a primary determinant of range carrying capacity in African savannahs, Shorrocks 2007), which would have been most interesting and required for any confidence in their conclusion that carrying capacity was not reached.

For SKKR, though IBI did not show any trends, AFR did increase with population size, even though this density dependence had no discernible effect on population growth (Law et al. 2013, 2105). As van Lieverloo et al. (2009) presented evidence against resource limitation for the SKKR population, Law et al. (2013, 2015) suggested the increase in AFR with population size was socially mediated. Greaver et al. also interpreted the density dependence evident in their regression as being socially mediated, presumably through the mortalities that generated the annual negative growth rates. We agree that density dependence may act on demographic factors such as AFR and IBI through social factors before it acts through resource limitation, especially in good quality black rhinoceros habitat, but we are not convinced that Greaver et al. have ruled out any resource limitation just on the basis of mean values of AFR and IBI. Possible trends in these demographic parameters need to be investigated.

Nevertheless, Greaver et al.’s regression is indeed intriguing but in our interpretation indicative of a carrying capacity being reached for the Ithala population, either purely social or possibly with a component of resource limitation. More research on black rhinoceros populations is required to understand the nature of density dependence, both the form in which it manifests itself on annual growth rates and in the underlying factors controlling it. In this regard, it does appear that there might be more information that could be gleaned from the Ithala population. That social factors may determine a carrying capacity below that expected purely on the basis of nutritional resources is an issue requiring further research and careful consideration with any dataset.

### Discrepancies in Greaver et al.’s Data Presentation

To confirm our interpretation of Greaver et al’s Fig. 2b was plausible, we examined the data presented by Greaver et al. carefully. For the population counts for the study years 1990 – 2008, we are told (p. 439) the population sizes were 46 in 1990 and 43 in 2008, with data for the intervening years displayed in their Fig. 2a and Fig. 4. The tick marks and year labels on the horizontal axis in Fig. 2a are not particularly helpful, but one can label the 18 data points for the years of interest from right to left by counting back from 2008 to 1990, as done in our annotated copy. Inspection then shows that their Fig. 2a and Fig. 4 are inconsistent. For example, for 1994, the count in Fig. 4 appears to be 50 while the datum for 1994 in Fig. 2a is clearly below 50. On the other hand, the count for 1992 in Fig. 4 is clearly less than 50 while the datum for 1992 in Fig. 2a is above 50. There is agreement for 1990, 1996, 1998, 2006, and 2008, but the counts for 2000 and 2002 in Fig. 4 appear high compared to those in Fig. 2a, while the count for 2004 in Fig. 4 appears low compared to that in Fig. 2a. These discrepancies leave one wondering which data to trust when evaluating the analyses represented by Figs. 2b and 2c. In particular, what is one to make of the abundances plotted on the horizontal axis of Fig. 2b that appear not to represent the same abundance data plotted in Fig. 2a?

**Figure 4.**
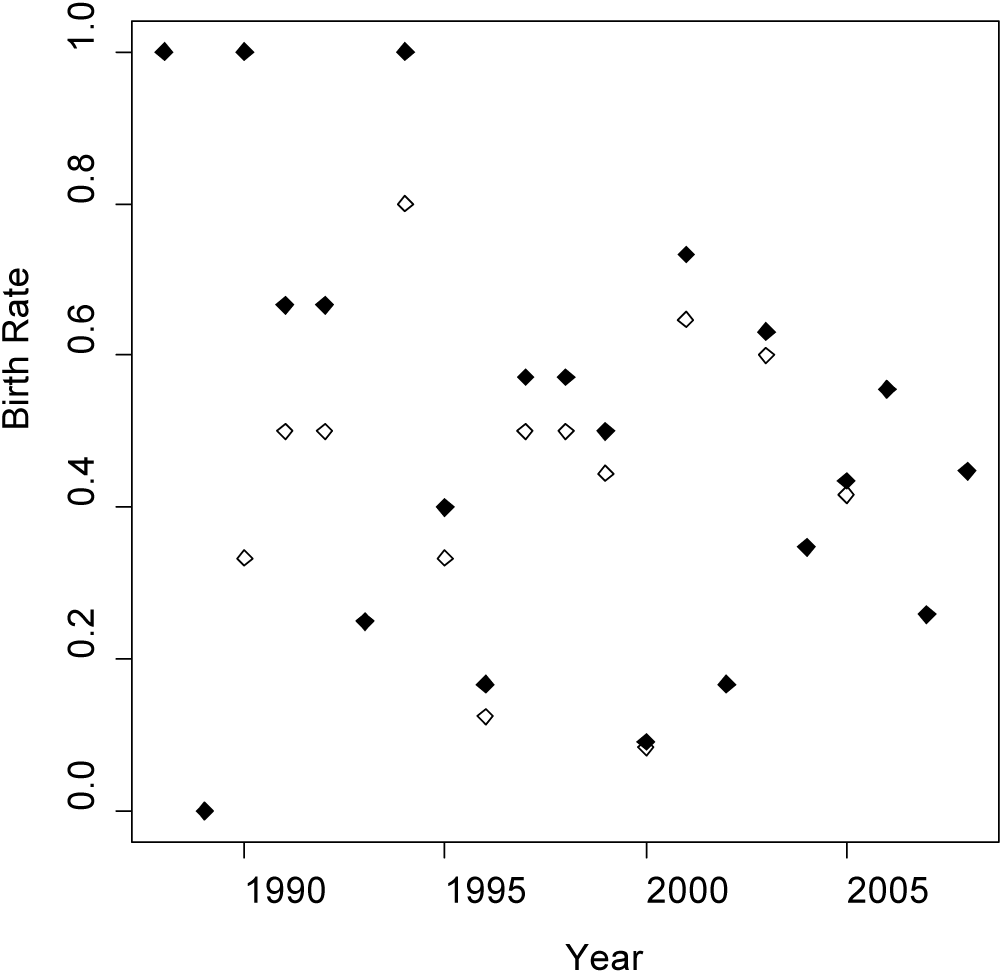
Annual birth rates (BR7 via Greaver et al.’s definition [closed diamonds]; BR counting all females that were at least seven years of age, as for BR7, plus any younger females that had already given birth [open diamonds]). Regressions for all data, post 1994, and post 1999 were all nonsignificant.

Greaver et al. computed the annual growth rate as *r* = ln(*N*_*t*+1_/*M*_*t*_), where *N*_*t*+1_ is the population size at the end of year *t* and *M*_*t*_ is the population size *N*_*t*_ at the beginning of year *t* minus the number of removals during that year (p. 439). Strictly speaking, our interpretation of their regression as a Ricker model assumes regression of *r* against *M*_*t*_. The annual removals are displayed in their Fig. 2a. Again, it proves useful to label these in decreasing order by year from right to left beginning with 2008. The meaning of the removals count for a year so labeled is still ambiguous. Is the removals count labeled 2008 for the year that ends with the 2008 count, i.e., these removals would be subtracted from the 2007 count in the computation of the growth rate for that year, or is it the removals for the year beginning with the 2008 count? First note that the 2006 count is the lowest count indicated by their Fig. 2a for 1990 – 2008 and, per their Fig. 4, is taken to be 37, which is consistent with their Fig. 2a. The removals for the year beginning with the 2006 count was nonzero, on either interpretation of the removal counts, and so the modified count corresponding to the 2006 count would be less than 37. Yet 37 is the smallest abundance displayed in their Fig. 2b. Thus, it appears that Greaver et al. have plotted, and regressed, *r* against *N*_*t*_ rather than *M*_*t*_, which means that our interpretation of the regression would be of a ramp model patching exponential growth with a modified Ricker model in which animals removed during the year are counted as contributing to density during that year. This fact does not undermine our proposed interpretation of the regression of their Fig. 2b as part of a ramp model.

We now assume that the abundances on the horizontal axis of their Fig. 2b are (unmodified) population counts at the beginning of the year for which the growth rates are computed. How do these counts accord with the inconsistent data in their Figs. 2a and 4? First note that their Fig. 2b has 18 distinct points plotted for the 18 annual growth rates, so there are no data hidden behind other points. In their Fig. 2a, the data for 1990 – 1997 are the highest abundances, with the remaining data in their Fig 2a below their level. So the data for 1990 – 1997 should be the eight highest values on the horizontal axis in their Fig 2b, apparently values of 53, 50, 49 (x 2), 48, 47 (x 2), but then there are two values of 46 on that axis, giving nine data not eight. (Note that the abundance for 2008 will not appear on the horizontal scale in their Fig. 2b.) Moreover, *three* of the points (1992, 1993, and 1997) in their Fig. 2a appear to be above the abundance level 50, not just one as in their Fig. 2b. Additionally, according to their Fig. 2b there is only one abundance less than 40, namely 37, but in their Fig. 2a there appear to be at least two, 2001 and 2006, while the counts for 1998, 2003, 2004, and 2008 are similar (the value for 2008 is 43 per p. 439). So where in their Fig. 2a is the 45 datum indicated in their Fig2b?

The data of their Fig. 4 appear to be more consistent with that displayed in their Fig. 2b, yielding counts of: 46 (1990, also per p. 439); 48 (1992); 50 (1994); 49 (1996); 42 (1998); 45 (2000); 43 (2002, the bar is the same height as for 2008, for which the abundance has been stated on p. 439 to be 43, yet there is only one plot of 43 in their Fig. 2b, so perhaps it should be 42?); 40 (2004); 37 (2006); and 43 (2008, per p. 439). Accepting these values, the count data in their Fig. 2a, at least, appears to be inaccurate. The most important point, however, is to determine whether the regression in their Fig. 2b can be regarded as reliable.

To this end, we return to the interpretation of the removal data displayed in their Fig. 2a. If the 1998 count is 42 (per their Fig. 4), and the removals labelled 1998 (the highest removals number, of about 12) occurred in the year ending with the 1998 count, then there are no removals in the year beginning with the 1998 count, the count for 1999 appears to be about 40 (here we must rely on Fig. 2a alas), and *r* ≈ ln(40/42) = −0.05, but (42, −0.05) is not a plotted point in their Fig. 2b. Alternatively, if the removals labelled 1998 occurred during the year beginning with the 1998 count, then instead *r* ≈ ln(40/30) = 0.29, and (42, 0.29) does approximate a point in their Fig. 2b. On this interpretation, for the year beginning with the 1999 count, one would also expect *r* ≈ ln(45/40) = 0.12 and (40,0.12) does approximate a plotted point. (Note that under any interpretation, the removal of 12 animals in the year prior to or following the 1998 count would yield a modified count of about 30 and there is no such abundance plotted in their Fig. 2b, further confirming that it is unmodified abundances plotted in their Fig. 2b.) Since the 1997 count is the highest count in their Fig. 2a, assume it is 53 per their Fig. 2b. If the removals labeled 1998 are again for the year ending with the 1998 count, then *r* = ln(42/41) = 0.02, whereas their Fig. 2b shows an *r* of about −0.25 for the abundance of 53. If, on the other hand, the 12 removals occurred in the year beginning with the 1998 count, then for the year beginning with the 1997 count *r* = ln(42/53) = −0.23, there being no removals under this interpretation for the year beginning with the 1997 count. The point (53, −0.23) is consistent with their Fig. 2b.

Hence, to make sense of the regression in their Fig. 2b and rescue our interpretation of it as part of a ramp model of density dependence, we made the following assumptions. We : gave preference to the population counts presented in Fig. 4 when they disagreed with Fig. 2a; interpreted the removals labelled by us as year *n* as the removals occurring in the year that began with the count for year *n*; and regarded the abundances on the horizontal axis in Fig. 2b as the unmodified counts at the beginning of the year for which *r* is computed. Our interpretation of the regression has the implication that the Ithala population did reach a carrying capacity during the study period.

### Problems with Birth Rates

Unfortunately, our interpretation of the data to rescue their Fig. 2b creates problems for the data presented in their Fig. 2c. Greaver et al. computed birth rate as the ratio of the number of births during a year divided by the number of adult females (females of age at least seven years old) alive at the end of that year (p. 439). Since they plot a birth rate for 2008, that datum is presumably for the birth rate of the year ending with the 2008 count, their last count, or they had knowledge of the number adult females alive one year after the 2008 count (they in fact plot a birth rate for each year 1990 through 2008). Now, under our interpretation, about 12 animals were removed for the year beginning with the 1998 count of 42. Taking as above the 1999 count to be about 40, about 10 animals were recruited by birth. For the 1998 count, there were, from their Fig. 4, about 18 adult females and for the 2000 count about 19. As their Fig. 4 does not suggest any abrupt changes in the number of adult females from year to year, we assume there were at most 18 or 19 adult females in the 1999 count whence the birth rate for the year ending with the 1999 count as computed by Greaver et al. (p. 439) would be at least 10/19 = 0.53 or 10/18 = 0.56. In Fig. 2c, the birth rate plotted for 1999 is less than 0.2, however (and, lest we have misinterpreted the horizontal-axis labelling, even lower for 1998). To reduce our estimate by over half due to our ignorance of the actual number of adult females in the 1999 count would require more than doubling the estimate of 18 or 19 adult females, which is highly unlikely, especially given the removals during the year starting with the 1998 count. For another example, consider the 2007 count, which appears to be level with that for 1998 in their Fig. 2a and just lower than the known 2008 count of 43, whence we assume it is 42. It would appear one rhino was removed during the year beginning with the 2007 count according to our interpretation of the removal data, whence *r* = ln(43/41) = 0.04. There is indeed a value of about 0.04 plotted above a count of 42 in their Fig. 2b. We are told there were 15 adult females in 2008 (which is also consistent with their Fig. 4), so with two recruits in between the 2007 and 2008 counts, the birth rate is 2/15 = 0.13, again at odds with the point plotted at about 0.32 above 2008 (which we assume is the appropriate comparison, but even more at odds with the point plotted at more than 0.5 for 2007) in their Fig. 2c. For our final example, assume the *r* value of about 0.275 at abundance 37 in their Fig. 2b is for the year beginning with the 2006 count (apparently the lowest abundance in their Fig 2a and consistent with their Fig. 4). From their Fig. 2a, it appears about five or six animals were removed between the 2006 and 2007 counts and the 2007 count is again taken to be 42. Then *r* ≈ ln(42/32) ≈ 0.27 (five removals) or ln(42/31) ≈ 0.30 (six removals). The first figure is compatible with their Fig. 2b and recruitment was ten. From their Fig. 4 it appears there were 16 adult females in the 2006 count, 15 in the 2008 count, so probably either 15 or 16 in the 2007 count, whence the birth rate for 2007 (or possibly 2006) is 10/16 = 0.625 or 10/15 = 0.67, both too high for any values plotted in their Fig. 2c (a smaller count of adult females if some were removed during the year only increases our estimate; it is perhaps possible that the number of adult females in 2007 was higher; 19 would turn our estimate into 0.53, which would be consistent with their Fig. 2c in this case).

Note that we are computing the birth rate as the number of recruits during the year divided by the number of adult females at the end of that year, rather than the number of births as done by Greaver et al., as they did not present that data, so our estimate of birth rate would be an underestimate if any animals died during their first year. Greaver et al. reported eight ‘juvenile’ (< 1 year old) deaths. This fact does not explain the discrepancies for 1999/1998 and 2007/2006 since our estimates exceed their estiamtes, but could play a role in the discrepancy for 2008/2007, for which ours (0.13) was less than theirs (about 0.32 for 2008, 0.22 for 2007). Unfortunately, though there were many mortalities (48) during the 1990 – 2008 period of their study, Greaver et al. provided no explicit details on their distribution over time. Those mortalities must, however, explain the negative growth rates but that does not tell us when the juvenile deaths occurred. It would require about 15 × 0.32 ≈ 5 of the juvenile yearling deaths (62.5%, or 10.4% of all mortalities) to have occurred between the 2007 and 2008 counts to obtain the plotted birth rate of about 0.32 (if instead we should be referring to the birth rate plotted above 2007 even more yearling deaths are required). Greaver et al. stated that yearling mortality was mostly due to cold/exposure but nothing was said to indicate it was concentrated in any particular year.

Thus, while we have presented a consistent interpretation of their Figs. 2b and 4, consistent also with their Fig. 2a, except for some inaccuracies in the count plots in that figure, it is apparently at odds with the birth rates plotted in their Fig. 2c. Even if this discrepancy is somehow due to our oversight, one might have expected greater scatter in the annual birth rates than is shown in their Fig. 2c. Our Figure 4 shows the plot for the SKKR population. Two datasets are plotted: annual birth rates as computed by Greaver et al., coded as BR7, i.e., the number of calves born during the year divided by the number of females of age at least seven years; alternatively, we counted as adult female all female rhinos at least of age seven, as Greaver et al. did, plus any younger females that had already given birth, and coded the resulting birth rate as BR (so Greaver et al. overestimated actual birth rate). Regressions of both the BR7 and BR data for SKKR, for all years, the years post 1994, and the years post 1999, were all nonsignificant, i.e., restrictions to periods of time excluding the earliest years of the reintroduction also failed to yield a significant regression.

The mean annual birth rate reported by Greaver et al. was 0.12. Note that the reciprocal of their mean IBI is 0.31, which should give a rough estimate of mean calving rate. For SKKR, the mean of annual birth rates as computed by Greaver et al. was 0.498, the mean using all known adult females (not just those aged at least seven as done by Greaver et al.) was 0.420, a direct computation of the number of births divided by the total number of adult female years during the SKKR study (1986 to 2008) yielded 0.440, while the reciprocal of the mean of all IBIs was 0.414. Thus, using the Greaver et al method for computing annual birth rate results in the largest estimate of mean annual birth rate, while the two means (of BR7 and BR) bracketed the direct computation and the reciprocal of IBI is quite close to mean(BR). Nevertheless, the ratio of mean(BR7) to the reciprocal of mean(IBI) was 1.20 for SKKR but only 0.39 for Greaver et al.’s data.

Thus, we find Greaver et al.’s plotted birth rates perplexing and in any case strike us as odd due to the surprisingly orderly plot of their Fig. 2c, while their mean annual birth rate is apparently low given their mean(IBI), which is at least worthy of comment if correct.

### Annual Birth Rates for Particular Months

Greaver et al.’s Fig 3 (not reproduced as we do not question the data) is a histogram of births by calendar month for 1990 – 2008. They declared the four months with the highest counts to be ‘peak birth months’: Mar (12); Jul (11); Apr (8); and May (7); but note that Jun, Aug, and Oct each had six births so the status of Apr and May a peak birth months is questionable. Their Table 1 shows results for models of ‘birth rate’ for each peak month, with two rainfall measures (*R*_15_, the cumulative rainfall over the 15 months prior to birth; R_27_, the cumulative rainfall over the 27 months prior to birth) and two measures of density (*D*_*b*_, population density in the year of birth; D_*c*_, population density in the year of conception) as predictors. Unfortunately, clarification is necessary to understand their modelling exercise. As they modelled ‘birth rate’, we assume they computed for each instance of, say, March, 1990 – 2008, the number of births in that March divided by the number of adult females alive at the end of that year (consistent with their previous definition of birth rate) and performed linear regression of birth rate against the four predictors. The first point of concern is that for no calendar month is there much variation in the number of births over the years 1990 – 2008. Even for March, with 12 births over 19 years, few months are likely to have more than one birth. Indeed, Table 1 lists a parameter *n* that is the same for each model for a given peak birth month and is, presumably, the sample size for the model. This parameter is ten for Mar, five for Apr, six for May, and nine for Jul. Focusing on March, we assume then that there are just ten birth rates from the 19 instances of March. Why? We can only assume that Greaver et al. excluded instances of March with zero births (though their covariates can still be assigned to such months and, moreover, excluding observations of zero seems erroneous to us). If so, then there were ten instances of Mar with at least one birth, and since there were only 12 births altogether, of the ten observations at most two can be of more than one birth. Hence, for any calendar month, there was very little variation in the number of births per instance over 1990 – 2008 (typically either zero or one). When the counts are converted to birth rates and zero counts ignored, it would seem that the variation in the data across the years is mostly due to variation in the conversion factor, i.e., variation in the number of adult females, which will be correlated with the population size, whence density (a rough estimate from the data in their Fig. 4 is a correlation of 0.6). Thus, one might expect to obtain a significant regression of birth rate on population density measures, with a negative regression coefficient. (Leaving aside the issue of lack of variation in the birth count data, we would have preferred a Poisson regression of the actual count data against the chosen predictors, possibly using a zero-inflated model, or a negative binomial regression.) Furthermore, the sample sizes are too small for many of the models considered, not just the one noted, but erroneously referred to as having too few parameters, in the footnote to their Table 1.

Greaver et al. concluded that ‘births were mostly explained by population density at conception, and the lag effect from rainfall on birth rate’ (p. 440). Using ΔAICc = 2 as a cutoff, the results of Table 1 suggest the modelling results are somewhat less clear cut. The only models to satisfy this cut off are: for Mar, the single model *R*_15_ x *D*_*c*_ + *D*_*b*_; for Apr and May each of the four models with a single predictor; for Jul, the two models with either *R*_27_ or *D*_*c*_ as predictor. So, it is true that *D*_*c*_ plays a role for each peak birth month, but the results for Apr and May are not particularly compelling, especially given our interpretation of the modelling exercise.

We have further queries about the modelling. The only evidence for a good fit of a global model to the data in each case are *R*^2^ values reported for each model, but it is adjusted *R*^2^ values that are more appropriate (since *R*^2^ always increases with model complexity). The parameter *K* in Table 1 is said to be the number of (structural) parameters in each model, including the intercept and each explanatory variable. Yet *K* = 1 for models with a single predictor, and *K* = 3 for a model such as *R*_15_ x *D*_*c*_ + *D*_*b*_. It would appear the intercept has not been included after all, while if the latter model included separate terms for *R*_15_ and *D*_*c*_ as is normal in a regression with interaction, there should be five structural parameters in this model (including the intercept). Moreover, if this *K* is the *K* employed in the computation of AIC_c_, then it must also count the estimated residual variance *σ*^2^ as a parameter (Burnham and Anderson 2002:12). One is left wondering what value of *K* was indeed used in the computation of AIC_c_ and hence whether the AIC_c_ values reported are correct. Irrespective of these technical points, we doubt the variation in the count data is sufficient to justify any analysis. Moreover, Greaver et al. did not report even the signs of regression coefficients. Presumably, they were negative for density (confirmed in Discussion on p. 442) and positive for rainfall. The former is what we predicted on purely statistical grounds. Furthermore, if birth rate is density dependent, is not that inference at odds with their assertion that AFR and IBI data do not indicate any density dependence in the Ithala population (p. 445) and with the simple trend of birth rate over time in their Fig. 2c?

## Conclusion

Greaver et al. concluded that the Ithala population dynamics were constrained by density dependence (birth rates constrained by density) but that the mean AFR and IBI values did not indicate resource limitation. They took mortalities due to fighting to indicate social interactions constraining population growth, i.e., the density dependence constraint is socially mediated, resulting in both deaths and presumably reduced birth rates. These conclusions are important if valid.

We have proposed an interpretation of their Fig. 2b as indicating a density dependent decline, consistent with a ramp model of density dependence as proposed for megaherbivores (Fowler 1981, 1989; McCullough 1992, 1999; Emslie 2001), though it remains unclear as to how density dependence was mediated. We challenge their analyses of birth rate, not being able to find an interpretation of their Fig. 2c that is consistent with the remainder of their Fig. 2 and Fig. 4, while rejecting their analysis of birth rates for specific months due to lack of variation in the number of births and questionable, or at least unclear, statistical procedures. Mean values of AFR and IBI alone are not sufficient to preclude trends with density, which if present may or may not be due to resource limitation or socially mediated behaviour. We believe the dynamics of the Ithala population require further clarification as regards the influences of density per se, how it is acting, and also the possible influence of environmental variation on demographic rates.

### Note

Our enquiries to the authors of Greaver et al, initially regarding some of the issues we raise in this paper, and subsequently to request their data or directions to the appropriate authority to obtain the data, went unanswered.

